# Controlled reassociation of multistranded, polycrossover DNA molecules into double helices

**DOI:** 10.1101/2025.08.22.671170

**Authors:** Nada Kabbara, Lauren Anderson, Arun Richard Chandrasekaran

## Abstract

Shape-changing DNA nanostructures have found applications in biosensing, drug delivery, cell modulation and data storage. A key aspect of this reconfiguration is the interaction of DNA nanostructures with other biomolecules or chemical stimuli such as pH and ionic conditions. Sequence-based nanostructure reconfiguration is largely achieved by strand displacement which is based on single stranded toeholds. In this work, we use sequence and temperature-controlled reassociation of one type of a DNA nanostructure into another structure. We demonstrate this strategy using the paranemic crossover (PX) DNA, a four-stranded structure with two adjacent double helical domains connected by six strand crossovers. In the presence of an anti-PX structure that is composed of strands that are each complementary to those in PX DNA, the structures reassociate at specific temperatures to form duplexes. Using the denaturing agent formamide, we decreased the temperature required for this reassociation. We extend the strategy to other polycrossover DNA molecules such as a double crossover motif (2 crossovers) and a juxtaposed DNA motif (4 crossovers), showing controlled reassociation of different DNA motifs into duplexes. Our study highlights the potential for DNA motifs to function as switchable molecular systems, offering new insights for responsive DNA-based materials and devices.

---

Reconfigurable DNA nanostructures play a key role in applying dynamic DNA nanotechnology in fields such as diagnostics, drug delivery, molecular computation and reactive circuits.^1^ Typically, reconfiguration is achieved through environmental (temperature), physical (light), chemical (pH, ions) and biomolecular (DNA, proteins, antigens) stimuli. For biomolecular reactions, While toehold-based DNA strand displacement^2^ is an often-used strategy to reconfigure DNA nanostructures, several recent works have focused on creating toehold-less strand displacement using weaker DNA interactions,^3^ enzymes^4,5^ or sequence composition.^6^ In addition to the sequence-based affinity widely used in strand displacement reactions, we recently showed structure-based affinity as an alternate strategy to reconfigure one DNA structure into another.^7^ Combinations of environmental stimuli and sequence specificity could result in control over reactions where DNA nanostructure reconfiguration into a different type of structure yields largely different properties.

In addition to developments in reconfigurable DNA nanostructures, there has been considerable effort towards the minimalistic synthesis of DNA nanostructures and simpler assembly routes.^8^ Only one component strand can be designed to self-assemble into nanotubes,^9^ 1D and 2D arrays,^10^ and 3D crystals,^11^ while two component strands can assemble into size-defined nanoprisms.^12^ By specific sequence design, two individual strands of a DNA duplex can each be assembled into similar DNA nanostructures, indicating the ability to create multiple DNA nanostructures in the same solution with a starting material that is different in structure.^13^ Enzymatic digestion of component DNA or RNA strands in duplexes can also trigger the formation of DNA or RNA cubes from the undigested strands.^14^ While component strands of a duplex can reassemble into DNA nanostructures, the opposite strategy has also been developed, where DNA nanostructures can reassociate into duplexes. The Afonin group has led the development of this strategy, constructing DNA cubes,^15^ RNA cubes^15^ and RNA polygons^16^ that can interact with other nucleic acid cubes or polygons respectively to form duplexes. Reassociation of these structures into duplexes allowed the control over functionalities of the structures, such as combining two split aptamers or creating active siRNA molecules from inactive starting materials.^17–19^ In these developments, the DNA and RNA nanostructures were made of double helical edges, and the structures reassociate into double helices.

In this work, we explored whether DNA motifs that are multistranded and contain multiple crossovers can reassociate into DNA duplexes. We present evidence for reassociation of a model nanostructure, the paranemic crossover (PX) DNA motif, into duplexes, with strategies to reduce the temperature requirement for the reassociation through the use of a denaturing agent. PX DNA contains two adjacent double helical domains connected by six crossovers, and has been used in the creation of 1D^20^ and 2D arrays,^21^ in the cohesion of topologically closed molecules,^22^ and in creation of single stranded knots^23^ and origami.^24^ Further, the PX structure has been implied to have biological roles in homologous recombination, making it a good polycrossover model structure for demonstrating this strategy.^25^ We also show that reassociation is possible in other polycrossover DNA motifs with different number of crossovers, with their thermal stability dictating the reassociation temperatures.

To evaluate whether PX molecules can reassociate into duplexes, we chose a PX sequence we have used in our earlier study (**Fig. S1**).^26^ We designed an anti-PX structure that is assembled from reverse complements of each of the four strands in the PX molecule (**Fig. 1** and **Fig. S1**). That is, each strand in the PX (indicated by a, b, c, d in **Fig. 1**) has a fully complementary partner in the anti-PX (a*, b*, c* and d*, respectively). We set out to find out whether the component strands of the PX and anti-PX molecules reassociate to form duplexes. We first validated assembly of the PX and anti-PX complexes in tris-acetate-EDTA (TAE) buffer containing 12.5 mM Mg^2+^ using non-denaturing polyacrylamide gel electrophoresis (**Fig. 2a**). Both structures formed with near-efficient assembly yields. We also assembled the four individual duplexes and validated them on non-denaturing PAGE. The PX and anti-PX showed melting temperatures of 58.5 °C and 61 °C respectively, consistent with earlier reports,^27^ while the melting temperature of the duplexes ranged between 74 and 81.5 °C (**Fig. 2b** and **Fig. S2**).

**Figure 1.**
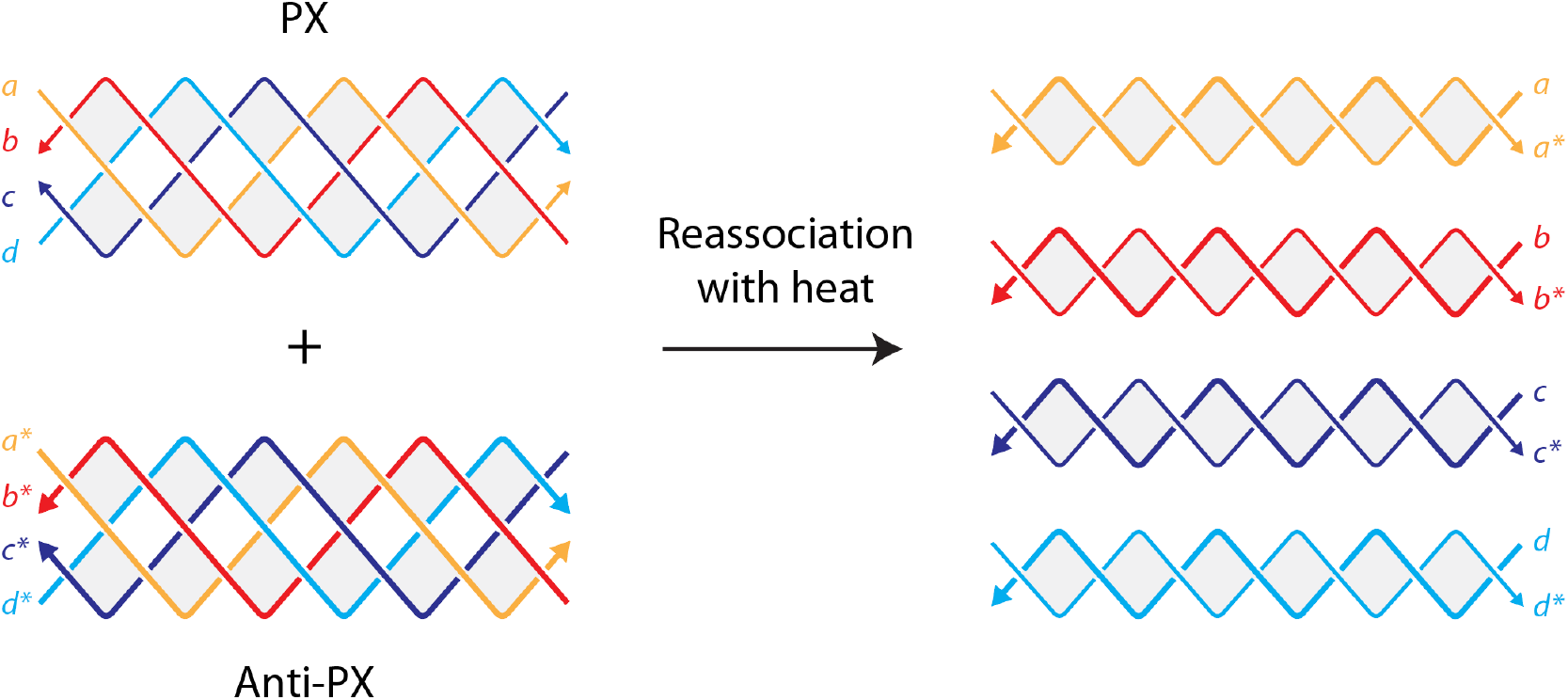
Reassociation of PX and anti-PX molecules into duplexes. The PX structure contains four strands hybridized into two adjacent double helical domains connected by six crossovers. Sequences of the strands in the PX are indicated by the letters a, b, c and d, each of which are 38 nucleotides long. The component strands of anti-PX molecule a*, b*, c*, d* are fully complementary to the component strands in the PX molecule. When mixed and incubated at specific temperatures, the complementary strands in the PX and the anti-PX molecules reassociate to form four duplexes a-a*, b-b*, c-c* and d-d*.

**Figure 2.**
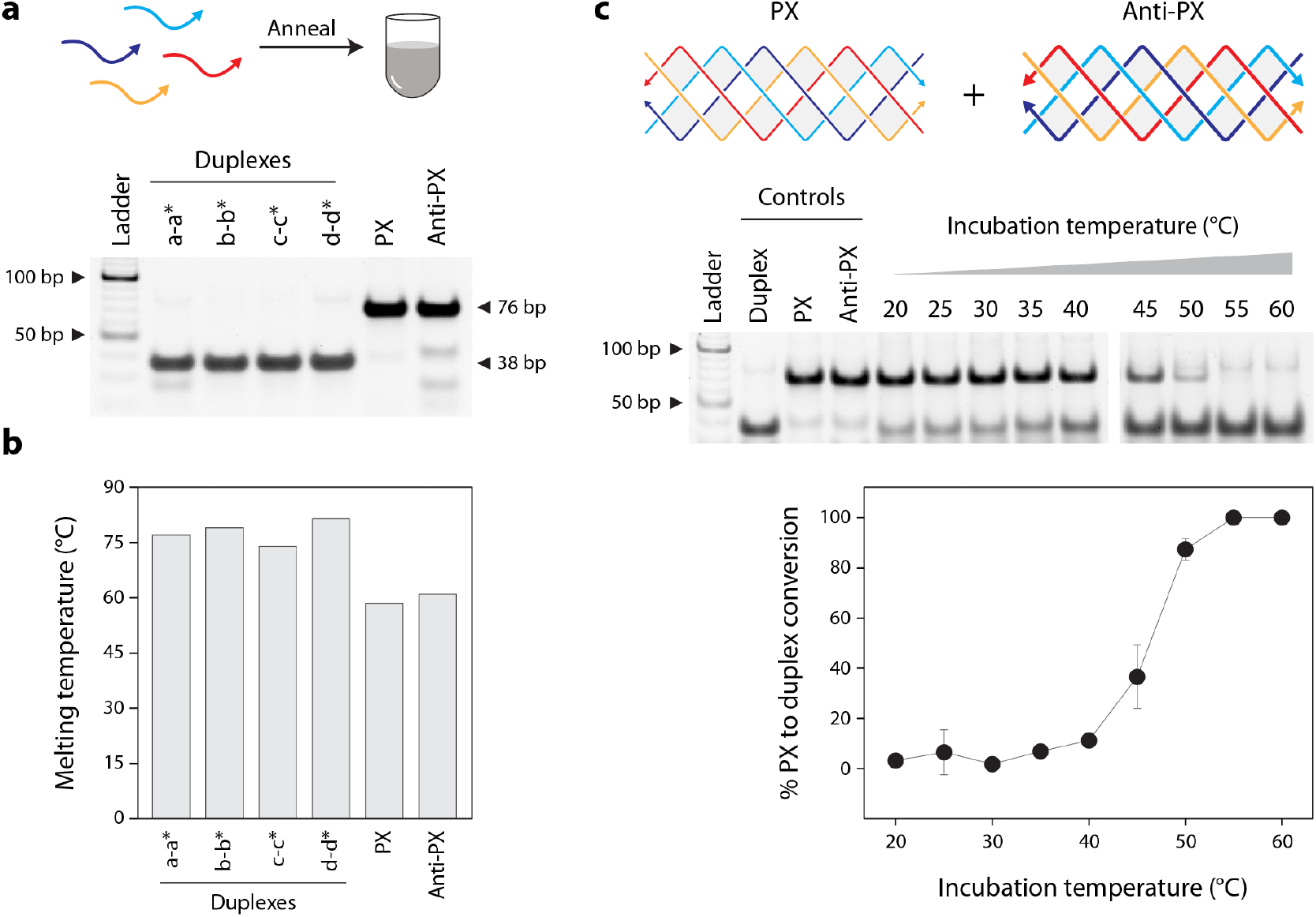
Assembly and reassociation of PX molecules. (a) Non-denaturing gel showing the assembly of the motifs and duplex structures used in this study. (b) UV melting temperatures of the structures. (c) Reassociation of PX and anti-PX molecules into duplexes at different temperatures. Reassociation is quantified as %conversion from PX to duplex based on the gel results.

Our initial notion was to assess whether the PX and anti-PX molecules reassociate at room temperature or physiological temperature (37 °C) to form duplexes if the individual strands in the PX preferred their complements in the anti-PX. However, we observed no reassociation at room temperature. Given the melting temperatures of the PX and anti-PX molecules, we then assessed whether the two structures could reassociate at higher temperatures. We assembled the PX and anti-PX, then incubated them together at temperatures ranging from 20 °C to 60 °C for 2 hours and analyzed the products on a non-denaturing gel (**Fig. 2c** and **Fig. S3**). We observed the reassociation of the PX and anti-PX into corresponding duplexes starting at a temperature of ∼40 °C, with the PX-to-duplex conversion increasing with temperature. At a temperature of 55-60 °C, all of the PX and anti-PX were reassociated into duplexes. To ascertain the stability of the PX and anti-PX at these incubation temperatures, we also analyzed the PX and anti-PX incubated individually at these temperatures. While the structures started denaturing at the higher temperatures tested, they were not fully denatured into the component single strands (**Fig. S4**). In combination with the melting experiments, the gel results showed that the PX and anti-PX molecules started to denature at these temperatures, allowing the individual complements to reassociate into duplexes. We note that reassociation of the duplexes into the PX molecules does not occur at the temperatures we tested, possibly due to the higher melting temperature of the duplexes compared to the PX and anti-PX molecules. This is also consistent with prior knowledge that formation of a single four-arm junction^28^ or polycrossover structures^29^ from intact duplexes is unfavorable and that duplexes are preferred over the crossover structures.

To improve the reassociation of PX into duplexes at lower temperatures, we added different amounts of formamide to the solution. Formamide has been shown to reduce the DNA melting temperature and is used in isothermal assembly of DNA nanostructures.^30^ We added 10-40% formamide in the solution containing the PX and anti-PX and incubated the mixture at temperatures ranging from 20 to 60 °C (**Fig. 3a** and **Fig. S5**). We observed a trend where the temperature required for the conversion of PX to duplex reduced with increasing formamide percentages in the solution (**Fig. 3b**). Thus, changing the solution conditions allows the complete reassociation of PX molecules into duplexes at temperatures as low as 30 °C.

**Figure 3.**
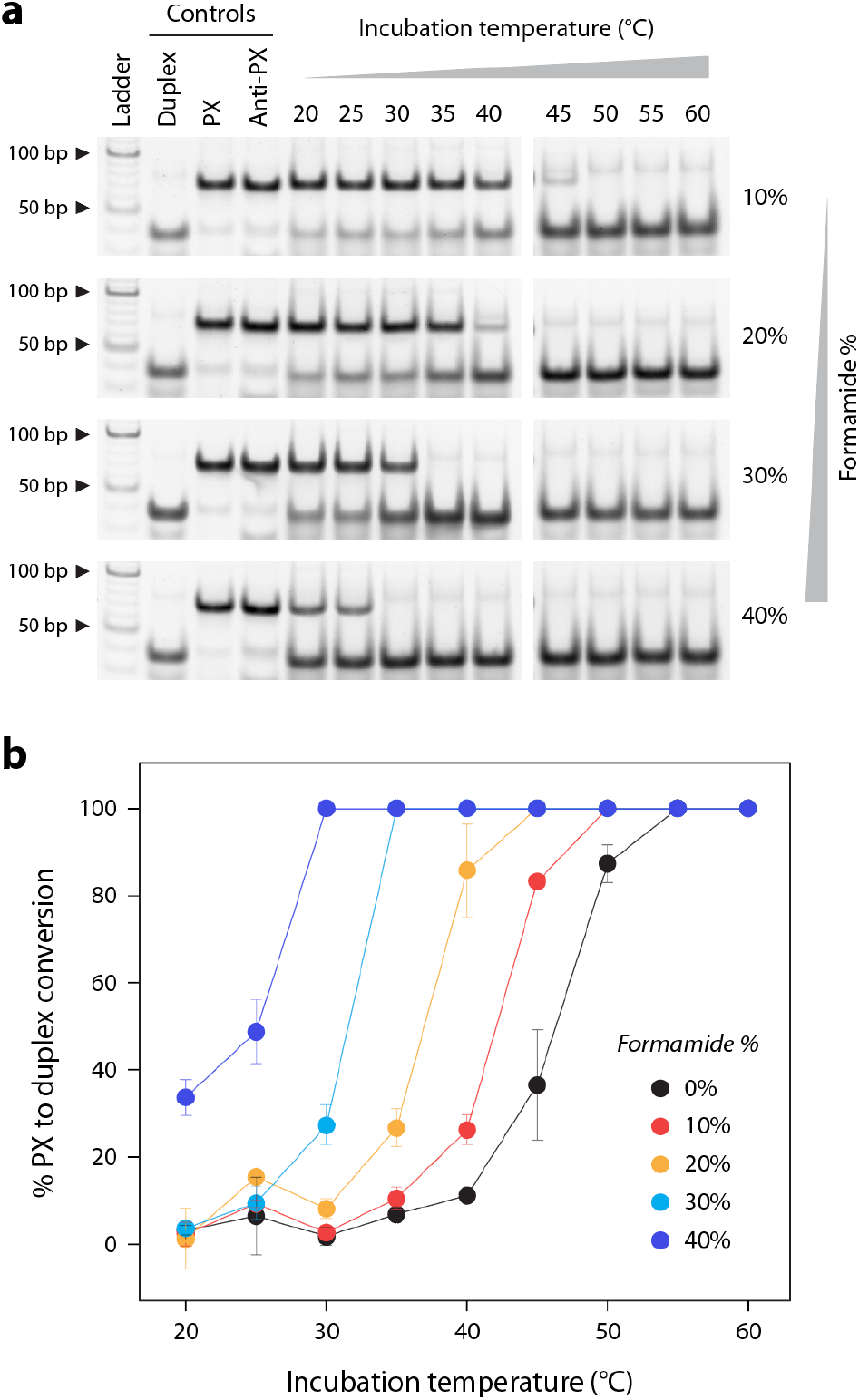
Decreasing reassociation temperature using denaturing agents. (a) Non-denaturing gels showing the reassociation of PX and anti-PX molecules into duplexes in the presence of different amounts of formamide. (b) Quantified results show a steady decrease in the temperature required for reassociation with increasing formamide concentrations.

Next, we used the reassociation of PX to duplex to demonstrate tunable biostability of DNA structures. Our earlier work showed that PX DNA has exceptional nuclease resistance when tested against nucleases and body fluids.^26^ Here, we incubated the PX/anti-PX mixture with different amounts of DNase I and observed minimal degradation over 30 minutes (**Fig. 4** and **Fig. S6**). We performed the DNase I assay at 20 °C so that there is no reassociation of the PX and anti-PX molecules, a temperature at which DNase I still shows activity. We then used the same starting batch of PX/anti-PX mixture and incubated it at 60 °C to ensure complete reassociation into duplexes. We incubated the reassociated duplexes with the same amounts of DNase I as before and observed that the duplexes degraded rapidly (**Fig. 4** and **Fig. S6**). We show that structures with high biostability (PX and anti-PX) can be reassociated into structures with weak biostability (duplexes), allowing one to introduce sequence-and temperature-based tunability between structures of vastly different biostability.

**Figure 4.**
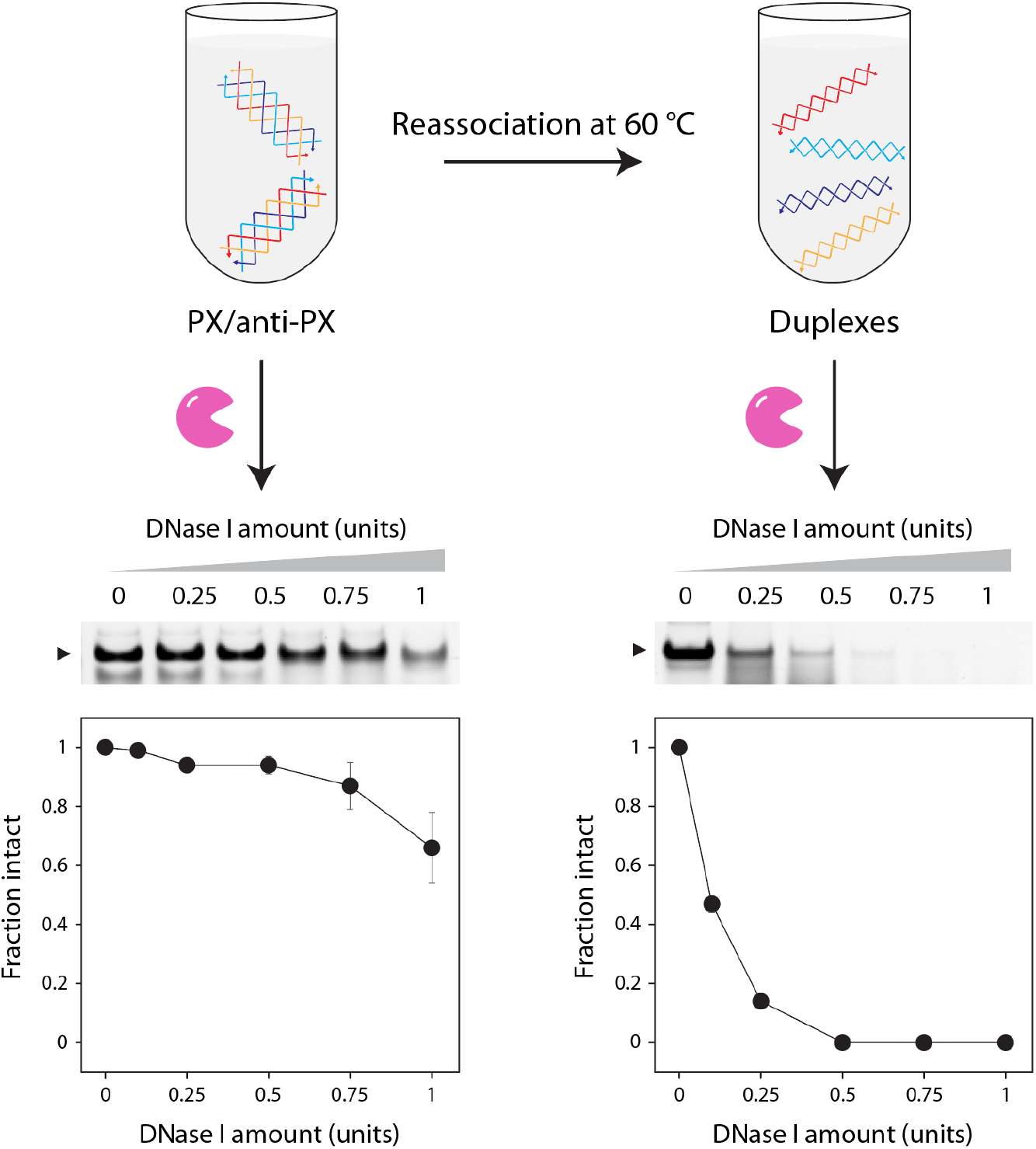
Tunable biostability. (Top) Illustration showing the DNase I treatment of PX/anti-PX mixture, and the reassociation of PX/anti-PX into duplex for DNase I treatment. (Bottom) Non-denaturing gel showing DNase I treated samples. PX/anti-PX molecules are nuclease resistant, showing only partial degradation over the range of nuclease concentrations tested, whereas the duplexes degrade completely.

Finally, we tested the effect of the number of crossovers in the structure on their reassociation temperature. We chose the double crossover (DX) motif that contains two crossovers and the juxtaposed crossover (JX) structure, a topoisomer of PX, that contains four crossovers (**Fig. 5a**). Similar to the PX and anti-PX, we designed anti-DX and anti-JX structures, where the component strands in anti-DX and anti-JX are fully complementary to the component strands in DX and JX, respectively (**Fig. S7-S8**). We incubated the DX/anti-DX and JX/anti-JX mixtures at different temperatures and observed that the two structures required different temperature ranges to reassociate into duplexes compared to the PX (**Fig. 5b-c** and **Fig. S9**). We found that the trend in the reassociation temperatures for these structures indicates a dependency on their thermal stability, rather than the number of crossovers in the structures. We observed that DX > PX > JX when comparing 50% reassociation into duplexes, a trend that is consistent with the melting temperatures of these structures (**Fig. S10**).

**Figure 5.**
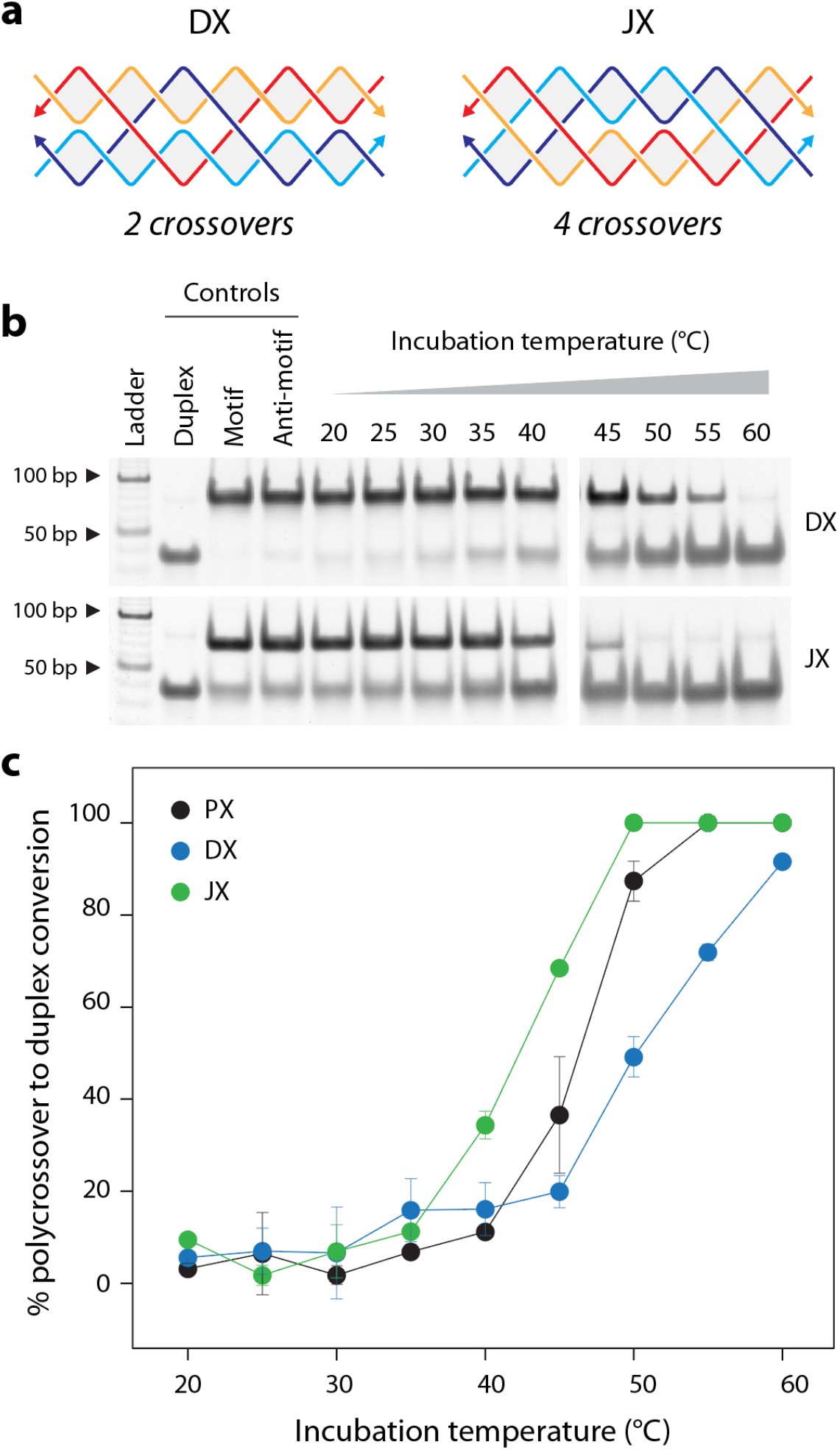
Reassociation of polycrossover DNA molecules into duplexes. (a) Scheme of a double crossover (DX) motif and a juxtaposed crossover (JX) motif containing four crossovers. (b) Non-denaturing gels and (c) quantified results showing the reassociation of the polycrossover molecules into duplexes at different temperatures.

In this work, we demonstrate that multistranded, tightly knit DNA motifs of a specific sequence composition can reassociate into duplexes. We show that reassociation can be achieved in a temperature-dependent manner, allowing the conversion of structures with vastly different nuclease resistance levels. While high temperature dependence is usually not preferred, operation of DNA nanostructures using heat as a stimuli is a useful feature, as shown in the cyclic transition of DNA origami dimers with fluctuating temperatures.^31^ For applications that require lower operating temperatures, we show that addition of formamide allows the reassociation temperature to go down by as much as ∼25 °C. Formamide has been used to enhance sensitivity of single molecule sensing using solid state nanopores,^32^ eliminate nonspecific interactions in gold nanoparticle-DNA sensors,^33^ accelerate the kinetics of DNA array transformation,^34^ in the folding of two individual strands of a double stranded DNA scaffold into independent structures^35^ and in the isothermal assembly of DNA polyhedra^36^ and origami.^37,38^ This strategy could also be combined with functional DNA modules that can assemble and reassociate in the presence of specific ligands.^39^ Since temperature is a key requirement, activation and reassociation can also be achieved by local heating as in nanoscale DNA heat engines that perform physical tasks such as moving beads on an origami.^40^

Structures such as the DX and PX DNA motifs are shown to be involved in biological processes such as homologous recombination.^41^ Thus our reassociation strategy has the potential to be adapted for biological applications where physiological temperatures can reassociate structures for different functionalities. Our prior work has also shown that PX and DX motifs do not cause any cell viability issues or adverse immune response, making these structures useful in biological applications.^26,42^ Similar to earlier work on using two strands of an siRNA that becomes functional after reassociation,^16,17^ our strategy could be useful for creating functional molecules after reassociation. Split aptamers and enzymes that become active once reassociation of the nanostructures combines the split pieces into duplexes might be useful in biosensing and drug delivery applications.^43^ In the context of dynamic DNA nanotechnology, our work indicates a route to create a group of nanostructures that can compute biomolecular information, resulting in reassociation into a different set of nanostructures, with properties that are useful in several biomedical applications.

## Supporting information

Experimental procedures, additional results and DNA sequences used.

## Author contributions

N.K. designed and performed experiments, analyzed data, and edited the paper. L.A. performed experiments and analyzed data. A.R.C. conceptualized and supervised the project, designed experiments, analyzed data, visualized data and wrote the paper.

## Acknowledgments

Research reported in this publication was supported by the National Institutes of Health (NIH) through National Institute of General Medical Sciences (NIGMS) under award number R35GM150672 to A.R.C. This manuscript is the result of funding in whole or in part by the National Institutes of Health (NIH). It is subject to the NIH Public Access Policy. Through acceptance of this federal funding, NIH has been given a right to make this manuscript publicly available in PubMed Central upon the Official Date of Publication, as defined by NIH. We thank Bharath Raj Madhanagopal for technical assistance with UV melting studies.

