## Supporting information for "Controlled reassociation of multistranded, polycrossover DNA molecules into double helices"

### MATERIALS AND METHODS

#### Preparation of DNA complexes

DNA strands were purchased from Integrated DNA Technologies (IDT). Full sequences are listed in Table S1. PX, DX and JX complexes were prepared by mixing the component DNA strands in equal ratios in Tris-Acetic-EDTA buffer containing 40 mM Tris base (pH 8), 20 mM acetic acid, 2 mM EDTA, and 12.5 mM magnesium acetate (1× TAE-$Mg^{2+}$). Samples were placed in a beaker containing 2 liters of deionized water heated to 90 °C, then placed in a Styrofoam box to cool to 20 °C over the course of 2 days. For duplexes, strands were combined in equal ratios in 1× TAE-$Mg^{2+}$ buffer and annealed using a thermocycler from 90 °C to 20 °C over 30 minutes.

#### Gel electrophoresis

Non-denaturing gels were prepared using 19:1 acrylamide/bisacrylamide (National Diagnostics). Samples were mixed with loading dye containing bromophenol blue and glycerol prior to loading. Gels were run at a constant voltage at 4 °C in 1× TAE-$Mg^{2+}$ running buffer. Gels were stained in 0.5× GelRed (Biotium), imaged using a Bio-Rad Gel Doc XR+ and analyzed using ImageLab software.

#### Structure reassociation

Assembled PX and anti-PX structures were mixed in equal molar ratios and incubated at different temperatures in a BioRad thermal cycler. The same procedure was followed for the DX and JX structures. Percent conversion from PX (or DX or JX) to duplex was calculated as the reduction in intensity of the band corresponding to multi-stranded structures (PX, DX or JX), and normalized to the anti-structure control band intensity.

#### Nuclease degradation assay

For PX samples, the PX and anti-PX were mixed at a final concentration of 0.5 µM and used for the DNase I assay. To convert the PX/anti-PX mixture into duplexes, the PX and anti-PX mixture was heated at 60 °C for 3 h and the resulting sample was used in the DNase I assay. DNA samples were first mixed with DNase I reaction buffer (provided by the vendor) to a final 1× concentration. Enzyme dilutions were made in nuclease-free water. For the nuclease degradation assay, 1 µl of the enzyme was added to 10 µl of the DNA sample containing the reaction buffer and incubated at 20 °C for 30 min. Incubated samples were mixed with gel loading dye containing bromophenol blue and 1× TAE-$Mg^{2+}$ buffer and run on non-denaturing gels to analyze degradation. Degradation profiles were obtained by normalizing the band intensity corresponding to the structure (at each enzyme concentration) to the control DNA lane without any enzyme.

#### UV Melting

UV melting experiments were performed on a Cary 3500 UV-Visible Spectrophotometer (Agilent) using 1 µM DNA concentration. Absorbance at 260 nm was recorded while samples were heated from 15 °C to 95 °C at a rate of 0.5 °C/min.

| Strand name | Sequence |
|---|---|
| PX1 | GTGGTATCATCAATGCTATGTGTAGGCTTAGACCTGAG |
| PX2 | ACTAGGTCGCAACAGACACAATACTTGACCGAATCACT |
| PX3 | AGTGAGTCTAACAAGTCACATATCTGTGATGATCTAGT |
| PX4 | CTCAGTTCGGTGCCTAATTGTGGCATTTGCGACACCAC |
| Anti-PX1 | CTCAGGTCTAAGCCTACACATAGCATTGATGATACCAC |
| Anti-PX2 | AGTGATTCGGTCAAGTATTGTGTCTGTTGCGACCTAGT |
| Anti-PX3 | ACTAGATCATCACAGATATGTGACTTGTTAGACTCACT |
| Anti-PX4 | GTGGTGTCGCAAATGCCACAATTAGGCACCGAACTGAG |
| JX1 | GTGGTATCATCAATGCCACAATACTTGACCGAATCACT |
| JX2 | ACTAGGTCGCAACAGATATGTGTAGGCTTAGACCTGAG |
| JX3 | AGTGAGTCTAACAAGTATTGTGGCATTTGCGACACCAC |
| JX4 | CTCAGTTCGGTGCCTACACATATCTGTGATGATCTAGT |
| Anti-JX1 | AGTGATTCGGTCAAGTATTGTGGCATTGATGATACCAC |
| Anti-JX2 | CTCAGGTCTAAGCCTACACATATCTGTTGCGACCTAGT |
| Anti-JX3 | GTGGTGTCGCAAATGCCACAATACTTGTTAGACTCACT |
| Anti-JX4 | ACTAGATCATCACAGATATGTGTAGGCACCGAACTGAG |
| DX1 | AGTGATTCGGTGCCTACACATATCTGTTGCGACACCAC |
| DX2 | GTGGTGTCGCAACAGACACAATACTTCACCGAATCACT |
| DX3 | ACTAGATCATCAATGCTATGTGTAGGGTTAGACCTGAG |
| DX4 | CTCAGGTCTAACAAGTATTGTGGCATTGATGATCTAGT |
| Anti-DX1 | GTGGTGTCGCAACAGATATGTGTAGGCACCGAATCACT |
| Anti-DX2 | AGTGATTCGGTGAAGTATTGTGTCTGTTGCGACACCAC |
| Anti-DX3 | CTCAGGTCTAACCCTACACATAGCATTGATGATCTAGT |
| Anti-DX4 | ACTAGATCATCAATGCCACAATACTTGTTAGACCTGAG |

**Table S1.** Sequences used in the study (written 5' to 3').

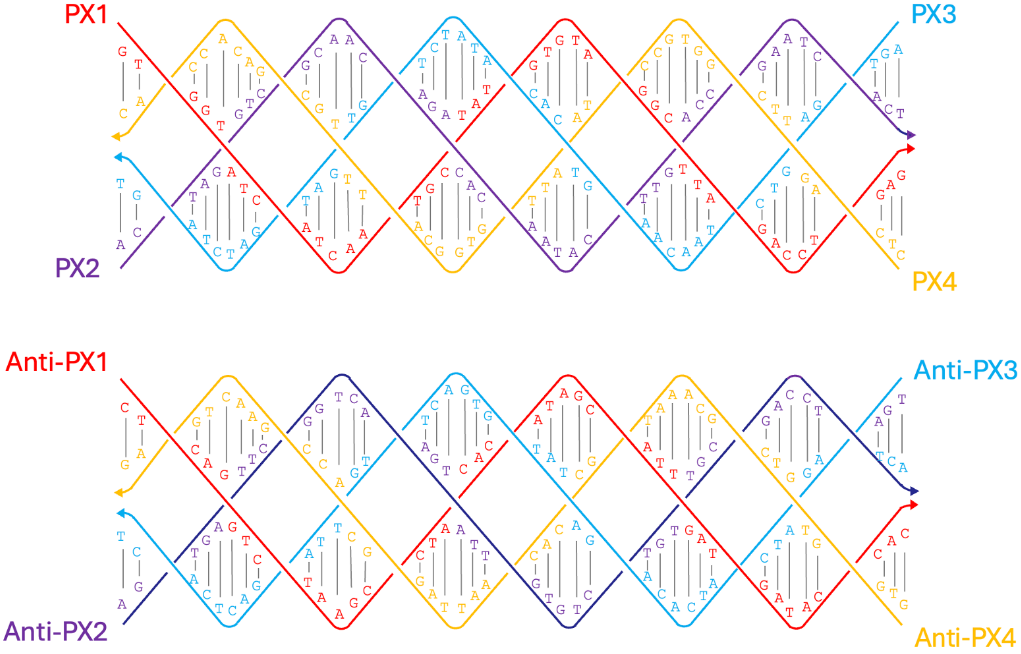


**Figure S1**. Sequence diagram for PX and anti-PX.

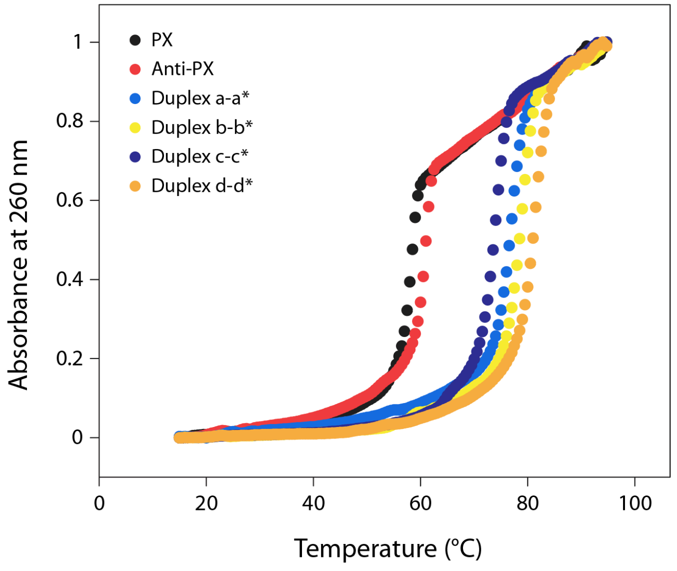


**Figure S2**. UV melting curves of PX and duplex molecules.

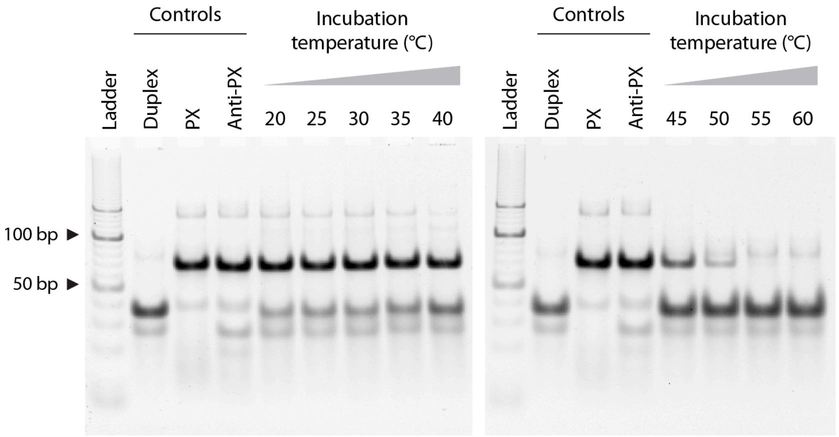


**Figure S3.** Non-denaturing gels showing the reassociation of PX and anti-PX in 1× TAE-$Mg^{2+}$ buffer at different temperatures.

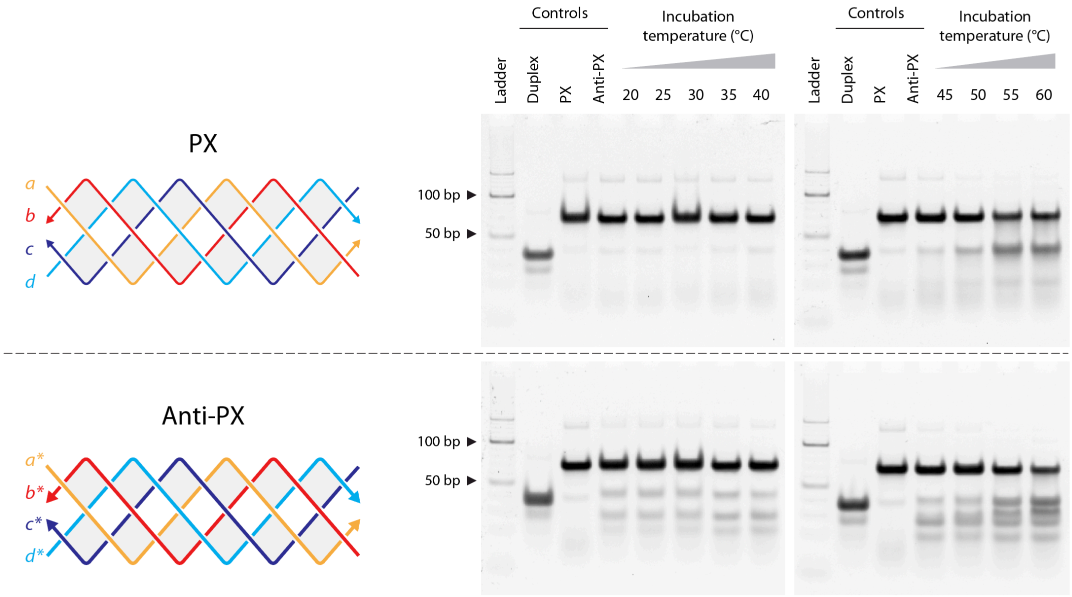


**Figure S4.** Non-denaturing gels showing stability of PX and anti-PX at different temperatures.

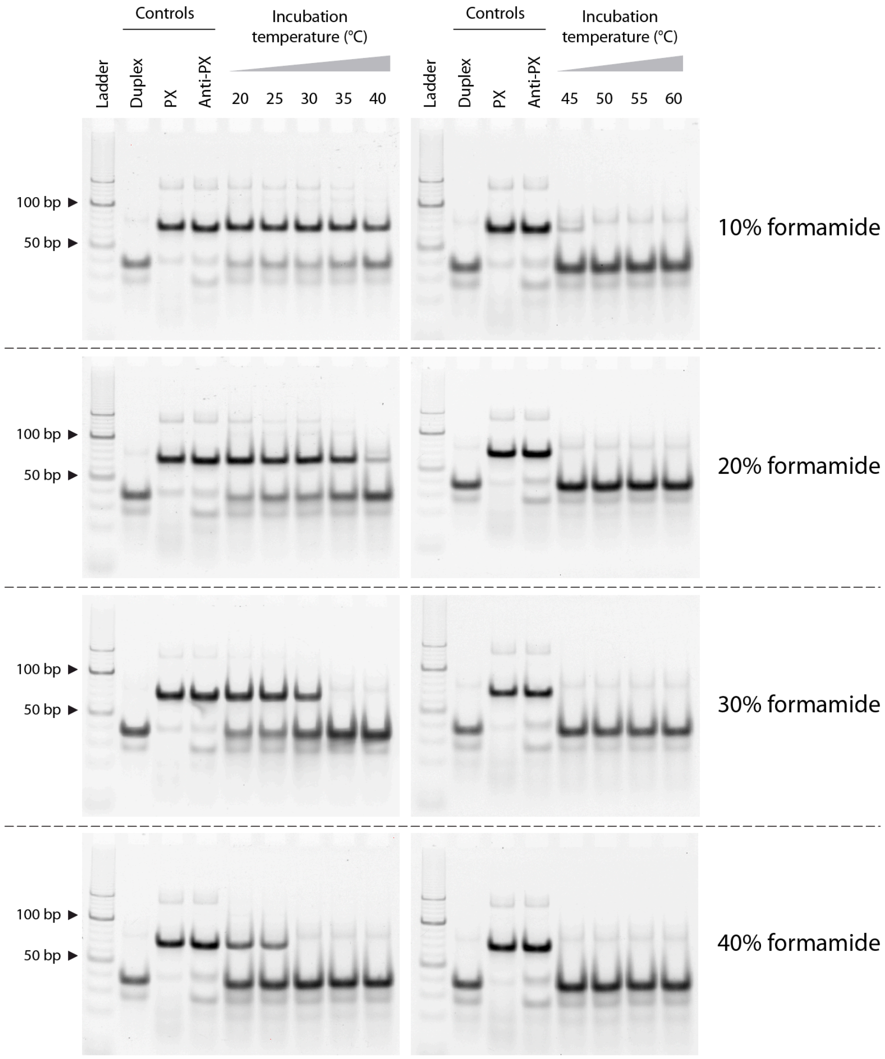


**Figure S5.** Non-denaturing gels showing reassociation of PX and anti-PX at different temperatures in 1× TAE-$Mg^{2+}$ buffer containing 10-40% formamide.

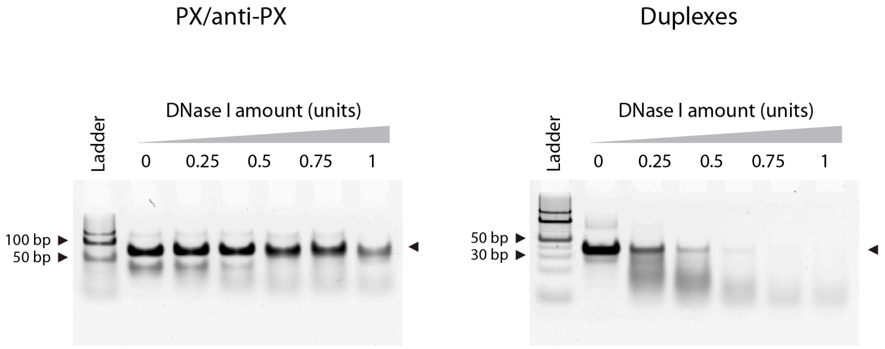


**Figure S6.** Non-denaturing gels showing DNase I degradation of PX/anti-PX samples (left) and reassociated duplexes (right).

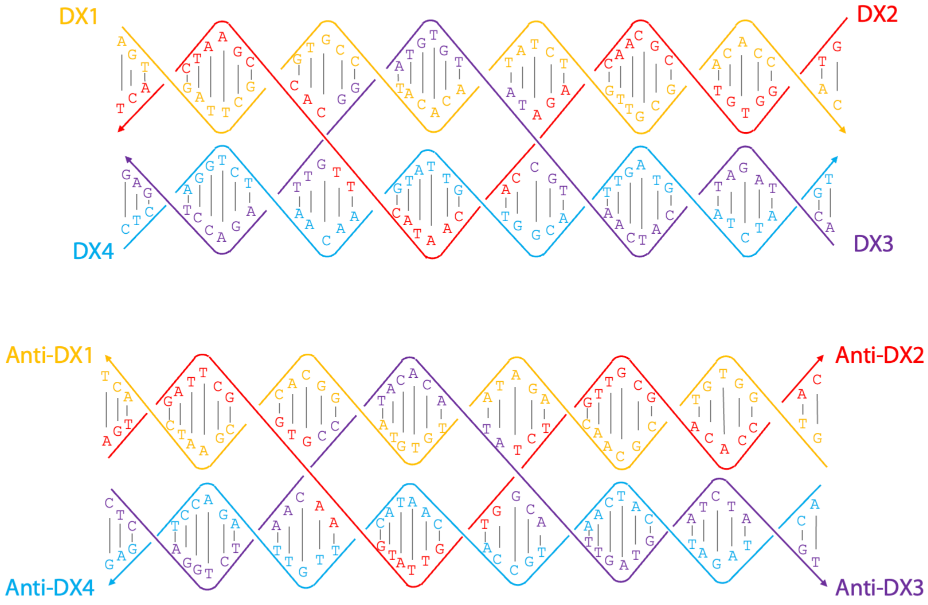


**Figure S7**. Sequence diagram for DX and anti-DX structures.

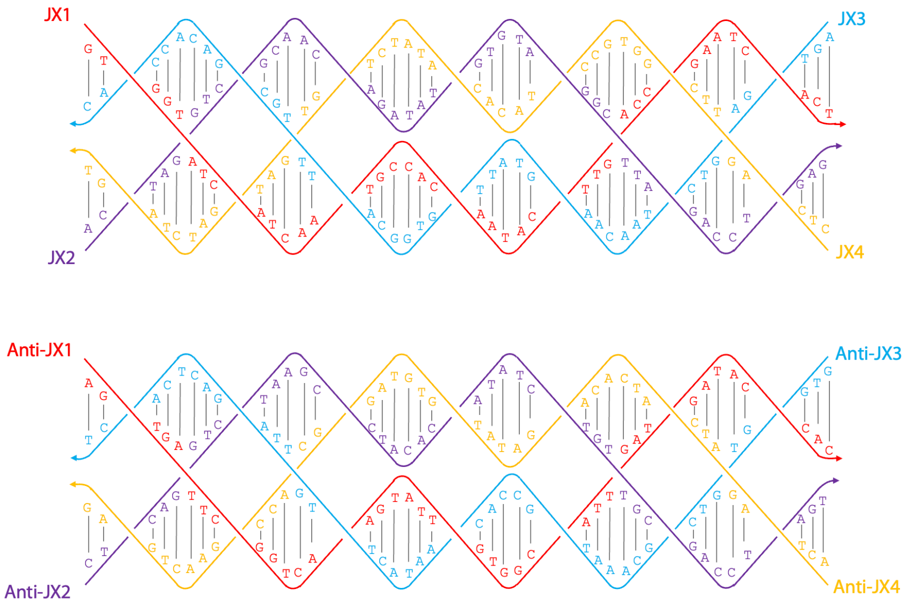


**Figure S8**. Sequence diagram for JX and anti-JX structures.

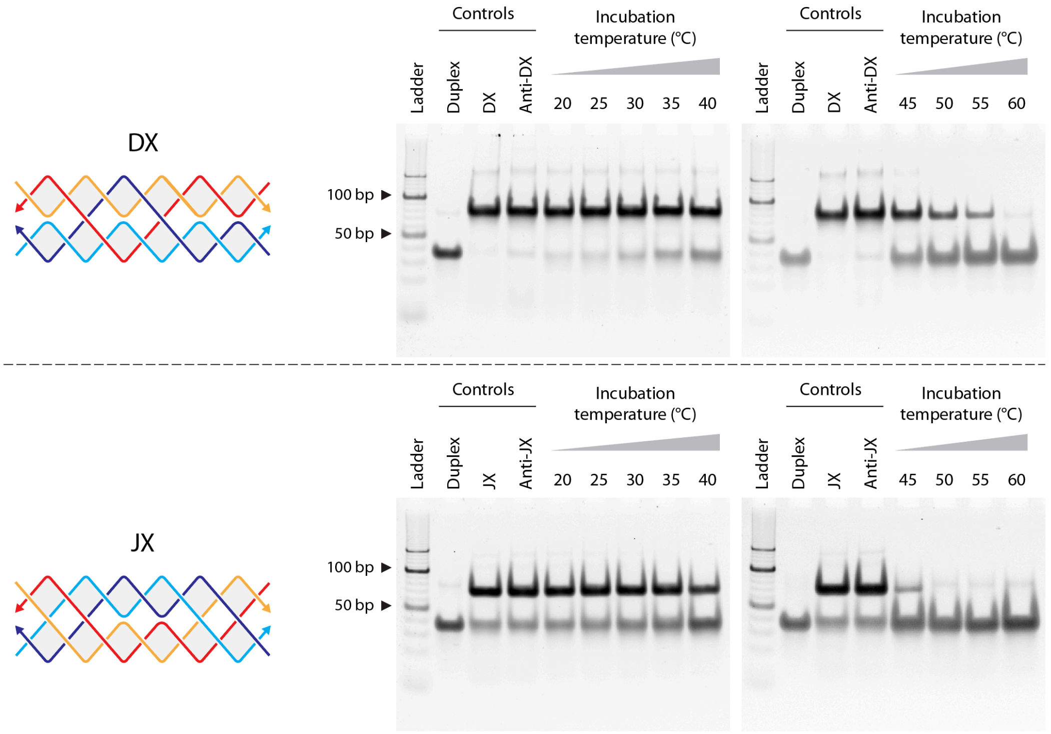


**Figure S9.** Non-denaturing gels showing the reassociation of DX and anti-DX (top) and JX and anti-JX (bottom) in 1× TAE-$Mg^{2+}$ buffer at different temperatures.

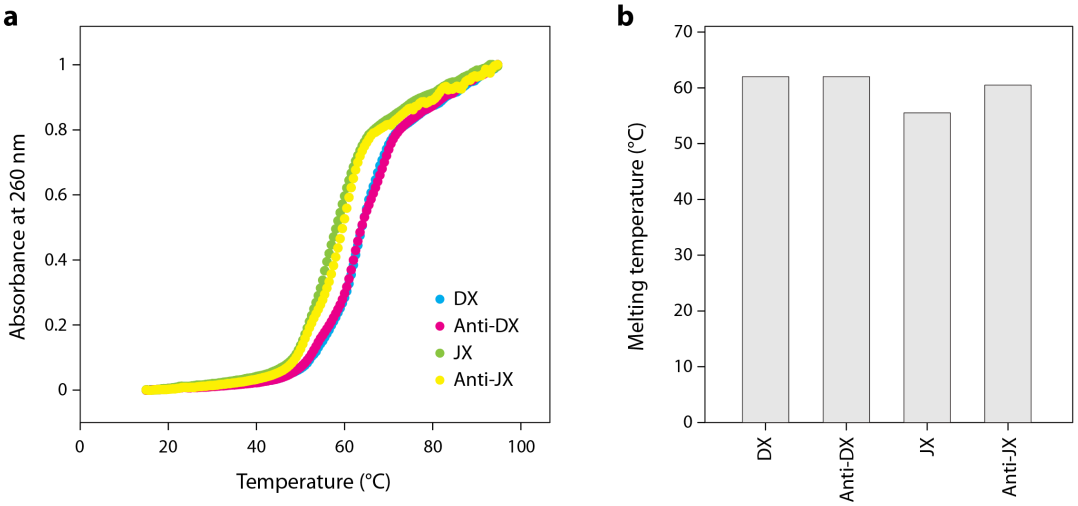


**Figure S10**. UV melting curves and melting temperatures for DX, JX and their corresponding anti-structures.
